# Brain-phenotype predictions can survive across diverse real-world data

**DOI:** 10.1101/2024.01.23.576916

**Authors:** Brendan D. Adkinson, Matthew Rosenblatt, Javid Dadashkarimi, Link Tejavibulya, Rongtao Jiang, Stephanie Noble, Dustin Scheinost

**Author notes:** Corresponding author: Brendan Adkinson.

## Abstract

Recent work suggests that machine learning models predicting psychiatric treatment outcomes based on clinical data may fail when applied to unharmonized samples. Neuroimaging predictive models offer the opportunity to incorporate neurobiological information, which may be more robust to dataset shifts. Yet, among the minority of neuroimaging studies that undertake any form of external validation, there is a notable lack of attention to generalization across dataset-specific idiosyncrasies. Research settings, by design, remove the between-site variations that real-world and, eventually, clinical applications demand. Here, we rigorously test the ability of a range of predictive models to generalize across three diverse, unharmonized samples: the Philadelphia Neurodevelopmental Cohort (n=1291), the Healthy Brain Network (n=1110), and the Human Connectome Project in Development (n=428). These datasets have high inter-dataset heterogeneity, encompassing substantial variations in age distribution, sex, racial and ethnic minority representation, recruitment geography, clinical symptom burdens, fMRI tasks, sequences, and behavioral measures. We demonstrate that reproducible and generalizable brain-behavior associations can be realized across diverse dataset features with sample sizes in the hundreds. Results indicate the potential of functional connectivity-based predictive models to be robust despite substantial inter-dataset variability. Notably, for the HCPD and HBN datasets, the best predictions were not from training and testing in the same dataset (i.e., cross-validation) but across datasets. This result suggests that training on diverse data may improve prediction in specific cases. Overall, this work provides a critical foundation for future work evaluating the generalizability of neuroimaging predictive models in real-world scenarios and clinical settings.

## INTRODUCTION

Machine learning offers the potential to augment clinical decision-making, individualize care, and improve patient outcomes (Johnson et al., 2021). Despite this promise, clinical neurosciences, particularly psychiatry, have yet to realize the advances in care that have been achieved by other medical disciplines. Recent work highlights that machine learning models predicting psychiatric treatment outcomes may be context-dependent and fail when applied to unharmonized samples (*i.e.*, across dataset shifts) (Chekroud et al., 2024). Given these models rely exclusively on clinical data, the addition of neurobiologically-grounded data, such as neuroimaging, may help overcome limitations due to inter-dataset variability (Sui et al., 2020).

In light of this, it is imperative to assess whether neuroimaging predictive models generalize across diverse dataset shifts. Only a minority of neuroimaging studies undertake any form of external validation. Among those that do, the median external sample size is only n=108 and is underpowered in most cases (Rosenblatt et al., 2023; Yeung et al., 2022). Further, real-world and eventual clinical applications demand not only external validation but also generalization across different imaging and phenotypic features (Dockès et al., 2021; Woo et al., 2017). By design, many consortium-level neuroimaging studies remove these variations, creating harmonization that does not exist in other scenarios. The inclusion of multiple datasets with different imaging parameters, patient demographics, and behavioral measures is necessary to truly evaluate a neuroimaging predictive model, as harmonization is not always possible (Chow et al., 2023; Torres-Espín and Ferguson, 2022). Models will only be clinically valuable if they can predict effectively on top of these dataset-specific idiosyncrasies.

In this work, we rigorously evaluate the external validation of neuroimaging predictive models across unharmonized samples (Figure 1). We use three distinct, large-scale developmental datasets: the Philadelphia Neurodevelopmental Cohort (PNC), the Healthy Brain Network (HBN), and the Human Connectome Project in Development (HCPD) (Alexander et al., 2017; Satterthwaite et al., 2016; Somerville et al., 2018). These datasets have high inter-dataset heterogeneity, encompassing substantial variations in participant characteristics (age distribution, sex, racial and ethnic minority representation, recruitment geography, clinical symptom burdens), imaging parameters (fMRI tasks and sequences), and behavioral measures. We used language abilities and function (EF) as two developmentally and clinically relevant phenotypes for prediction (Adise et al., 2023; Casey, 2023; Godfrey et al., 2022; Qi et al., 2021). We demonstrate that reproducible and generalizable brain-behavior associations using functional connectivity and connectome-based predictive modeling can be realized across diverse dataset features with sample sizes smaller than consortium-levels. Results indicate the potential of functional connectivity to be robust despite various dataset shifts. Further, they provide a critical foundation for future work evaluating the generalizability of brain-behavior associations in real-world scenarios and, eventually, clinical settings.

**Figure 1.**
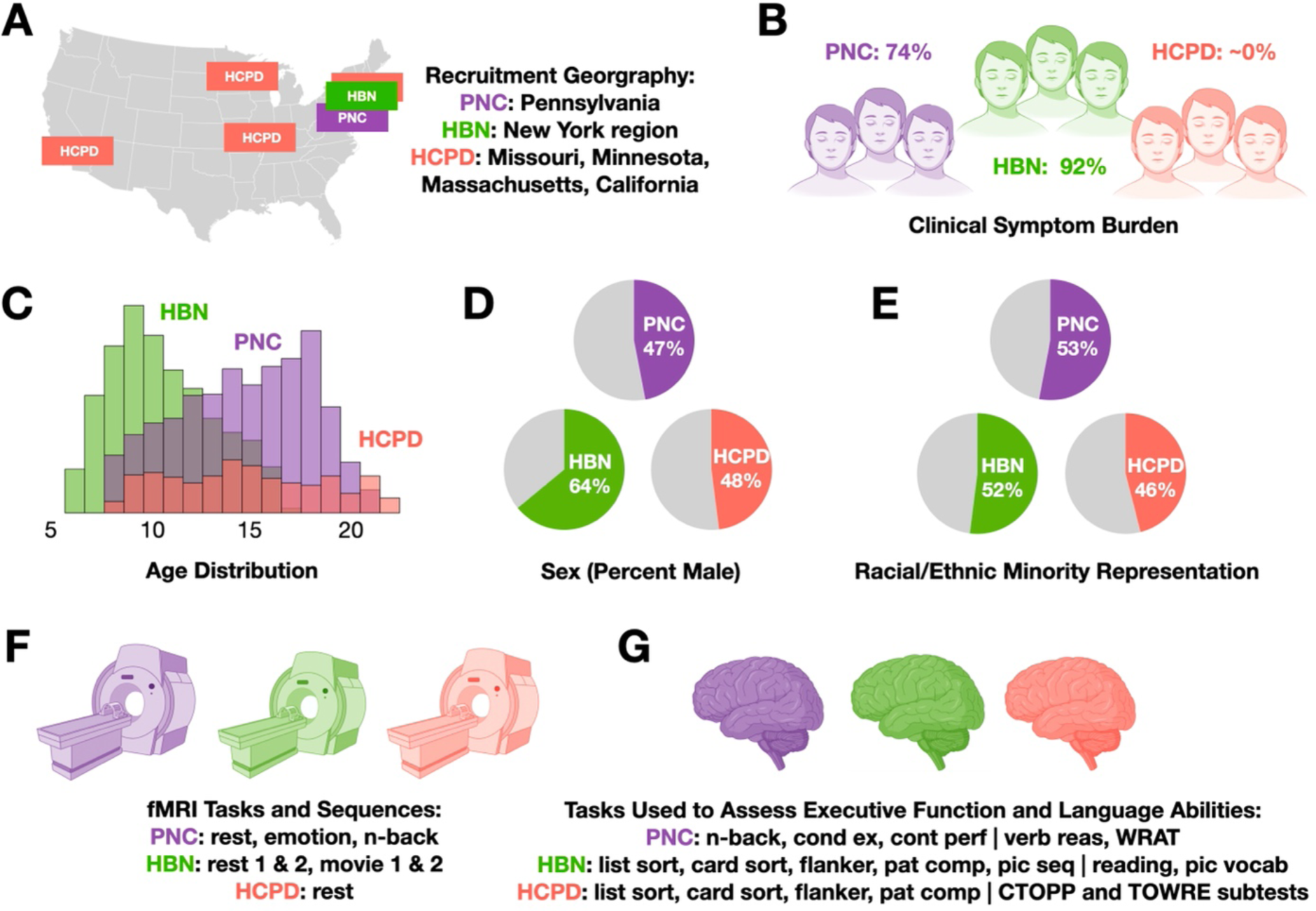
Differences across the PNC, HBN, and HCPD datasets. The Philadelphia Neurodevelopmental Cohort (PNC), Healthy Brain Network (HBN), and Human Connectome Project in Development (HCDP) datasets exhibit a notable lack of harmonization across recruitment geography (A), participant clinical symptom burden (B), age distribution (C), sex (D), racial and ethnic minority representation (E), fMRI tasks and sequences (F), and measures used to assess language abilities and executive function (G).

## RESULTS

We generated models of language abilities and EF in the PNC (n=1291), HBN (n=1110), and HCPD (n=428) datasets using ridge regression connectome-based predictive modeling (CPM) (Shen et al., 2017). Connectomes were created using the Shen 268 atlas. Each participant’s connectome included all available resting-state and task fMRI data with low motion (<0.2 mm). Combining connectomes across fMRI data improves reliability and predictive power (Elliott et al., 2019; Gao et al., 2019). Participants without one low-motion fMRI run were excluded.

A disparate set of behavioral tasks assessed language and EF in the three datasets (Table S1). We used principal component analysis (PCA) to derive “latent” factors of language abilities and EF within each dataset. Participants with missing language and EF measures were excluded. Importantly, the PCA was estimated using participants who did not have imaging data to maintain proper separation of training and testing data. The first principal component explained 70%, 55%, and 77% of language ability measure variance in PNC, HBN, and HCPD, respectively. For executive function, the first principal component of all behavioral measures explained 53%, 48%, and 40% of the variance in PNC, HBN, and HCPD, respectively. Contributions of individual measures to the first principal component are presented in Table S1. Behavioral data from participants with imaging data were projected onto the first principal component. This projection was used in all CPM analyses unless otherwise specified.

Predictive models were trained and tested within each dataset using 100 iterations of 10-fold cross-validation. Model performance was evaluated with Pearson’s correlation (r), representing the correspondence between predicted and observed behavioral scores, along with the cross-validation coefficient of determination (q^2^) and mean square error (MSE). Significance was assessed using permutation testing with 1000 iterations of randomly shuffled behavioral data labels. Cross-dataset predictions were evaluated with Pearson’s correlation.

### Connectome-based prediction of language abilities

Models successfully predicted language abilities within each dataset (Figures 2A and S1A; PNC: r=0.50, p<0.001, q2=0.24, MSE=1.05; HBN: r=0.27, p<0.001, q2=0.06, MSE=4.42; HCPD: r=0.22, p<0.001, q2=0.01, MSE=1.47). Model performance was similar to original predictions when controlling for age, sex, racial/ethnic minority representation, socioeconomic status, head motion, and clinical symptom burden (Table S2).

**Figure 2.**
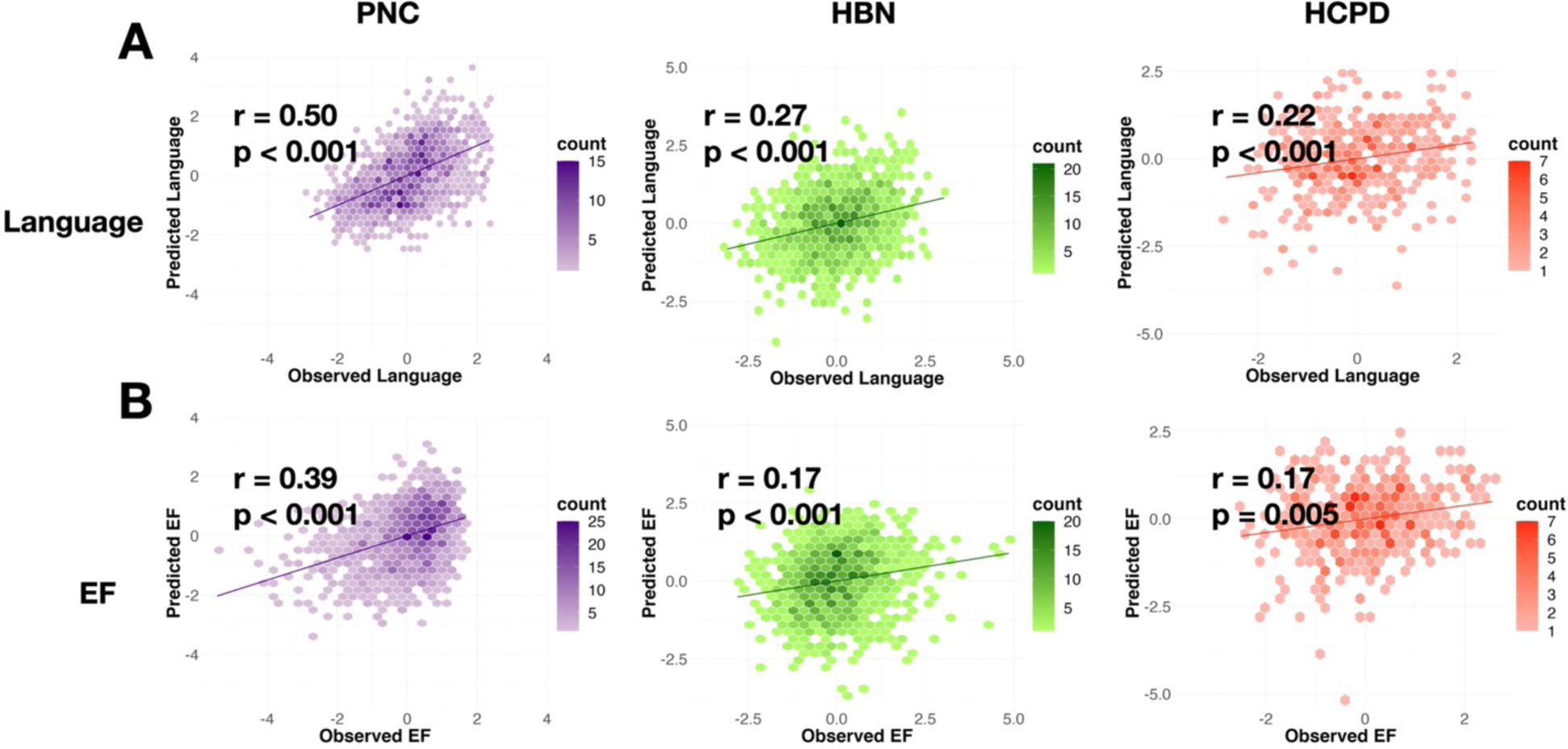
Connectome-based predictive model performance within-dataset. Scatter plot of observed 1st principal component scores on the x-axis and predicted 1st principal component scores on the y-axis for language abilities (A) and executive function (B) across PNC (purple), HBN (green), and HCPD (red). Counts represent individual participant data.

### Connectome-based prediction of executive function

The performance of EF models closely resembled the performance of language models (Figures 2B and S1B; PNC: r=0.39, p<0.001, q2=0.14, MSE=1.17; HBN: r=0.17, p<0.001, q2=0.02, MSE=2.03; HCPD: r=0.17, p=0.005, q2=-0.01, MSE=1.98). The addition of covariates into the model yielded similar results for age, sex, racial/ethnic minority representation, socioeconomic status, head motion, and clinical symptom burden (Table S2).

### Models generalize across datasets despite notable lack of harmonization

Cross-dataset predictions were performed across the three datasets to ensure our models’ generalizability. Importantly, PNC, HBN, and HCPD are characterized by a notable lack of inter-dataset harmonization (Figure 1). Despite such substantial differences, we achieved cross-dataset prediction of language abilities and EF (Figure 3). Language abilities were predicted with r’s=0.13-0.35. EF was predicted with r’s=0.14-0.28. Testing on the PNC produced the best cross-dataset predictions for language abilities and EF. As a result, the best predictions for the HCPD and HBN were not from training and testing in the same dataset (i.e., cross-validation).

**Figure 3.**
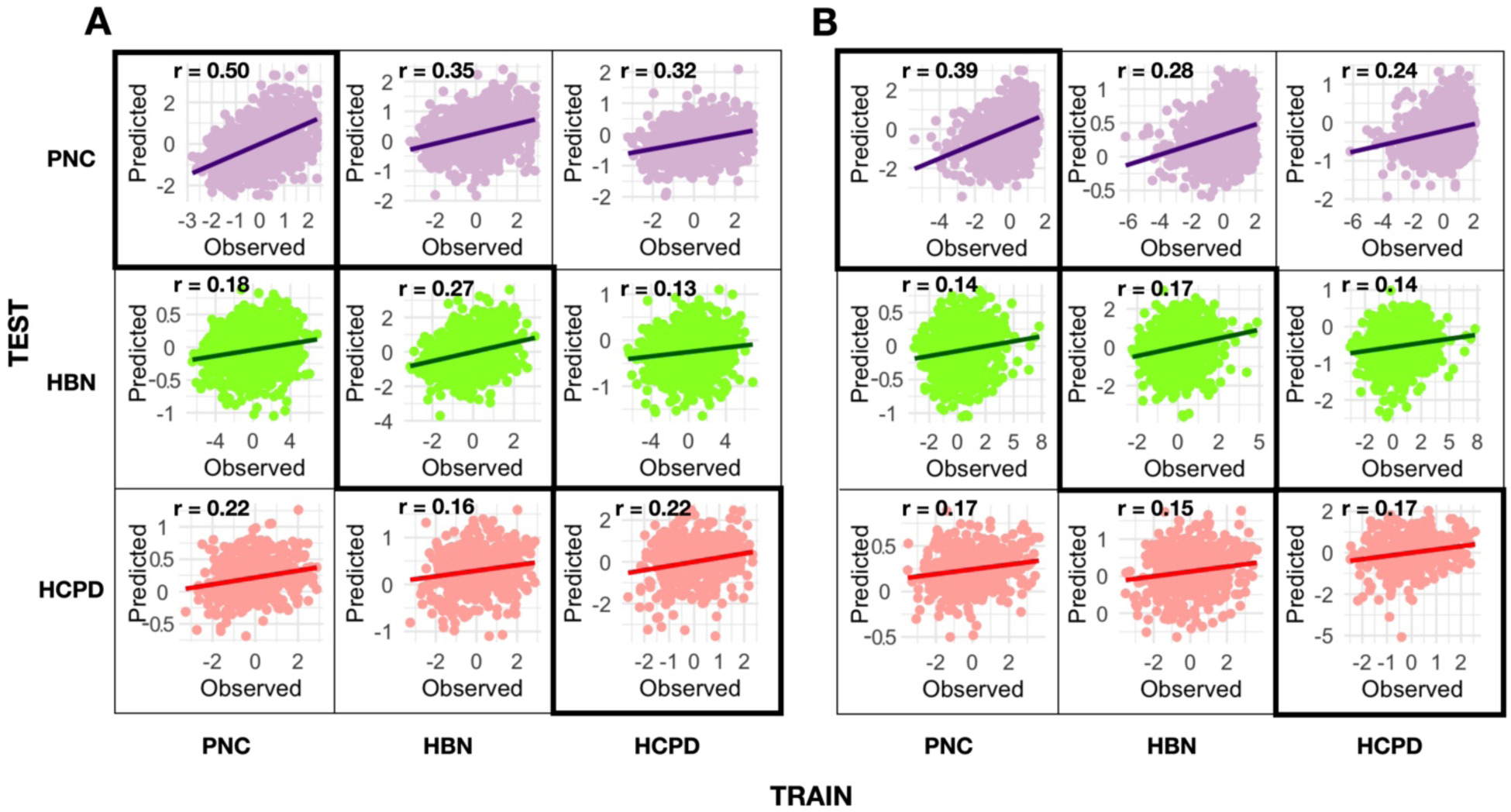
Model performances across unharmonized datasets. Scatter plots of true versus predicted PCA-derived language abilities (A) and executive function (B) scores for cross-dataset predictions. Purple (PNC), green (HBN), and red (HCPD) colors indicate the dataset in which predictions were tested. Diagonals represent within-dataset prediction performances.

### Brain features underlying language abilities and executive function

In line with previous CPM results, predictive models of language abilities and EF were complex, with contributions from every node and canonical brain network (Figures 4, S2). Virtual lesioning analyses confirmed the predictive utility of every brain network but also suggested the importance of the medial frontal and frontoparietal networks in predicting language abilities and EF (Figure S3). These networks contain noted regions for language (e.g., Broca’s and Wernicke’s) and EF (e.g., prefrontal cortex). We compared the brain features that predicted language abilities and EF in one dataset to those that predicted the same construct in the other two. All edgewise regression coefficients were normalized by the standard deviation of edges and summed for each canonical brain network. At the network level, predictive features from each dataset were correlated between r=0.48–0.74 for language abilities and r=-0.03–0.30 for EF. The correlations between the HCPD and the HBN or PNC were the lowest (Table S3).

**Figure 4.**
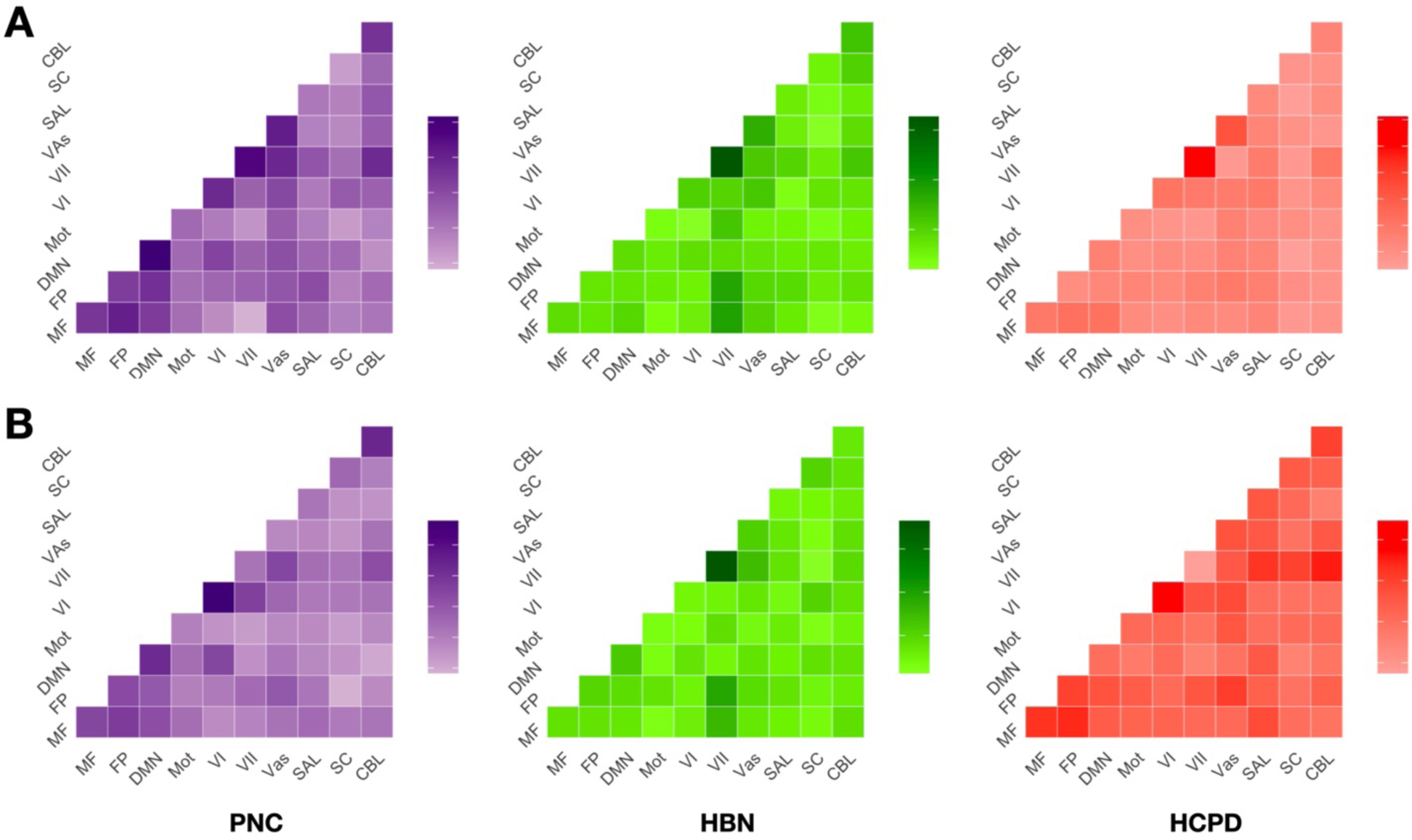
Network-level contributions to language abilities and executive function predictions. Canonical network contributions to predicted language abilities (A) and executive function (B) across PNC (purple), HBN (green), and HCPD (red). Contributions of edges within a single network (diagonals) and between networks (off-diagonals) were defined as the sum of edgewise regression coefficients normalized by network size. Darker colors indicate networks with larger model coefficients. Network Labels: MF, medial frontal; FP, frontoparietal; DMN, default mode; Mot, motor cortex; VI, visual A; VII, visual B; VAs, visual association; SAL, salience; SC, subcortical; CBL, cerebellum.

### Prediction of individual language and EF measures

Finally, we tested within and cross-dataset predictions for each measure used in the PCA. This analysis ensures that the strong cross-dataset predictions are not solely a function of combining disparate measures. Within-dataset predictions were significant across all individual measures, with the lowest being the HBN Card Sort task (r=0.07, p=0.05, Figure 5).

**Figure 5.**
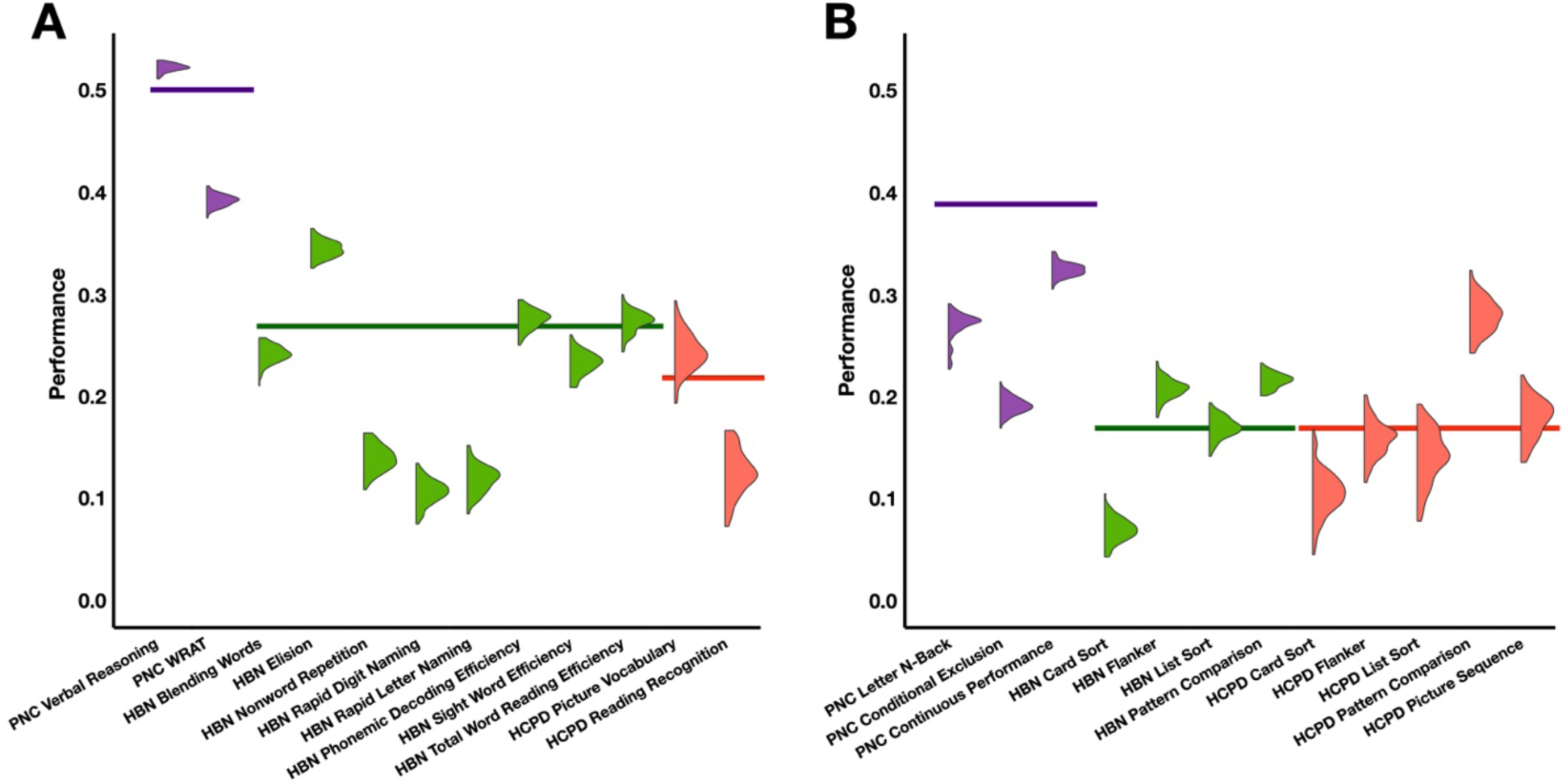
Within-dataset predictions of individual measures. Distributions of prediction performance Pearson’s r values across 100 iterations for each individual language (A) and EF (B) measure. PNC measures are purple, HBN measures are green, and HCPD measures are red. Solid lines indicate PCA prediction performances for comparison.

Cross-dataset predictions for the individual measures followed patterns similar to PCA-derived predictions and, in general, were significant (Figure 6). Mirroring PCA results, cross-dataset language abilities predictions (median r=0.14, interquartile range (IQR)=0.09) were more accurate than executive function predictions (median r=0.11, IQR=0.10). For language abilities, all individual measures were predicted in at least one cross-dataset model. 58 out of 72 cross-dataset models were significant, including all models tested in the PNC. For EF, 61 out of 94 cross-dataset models were significant. Models built on the flanker task showed the worst generalization. Most predictions used different measures in the training and testing data, showing strong generalization of language and EF models.

**Figure 6.**
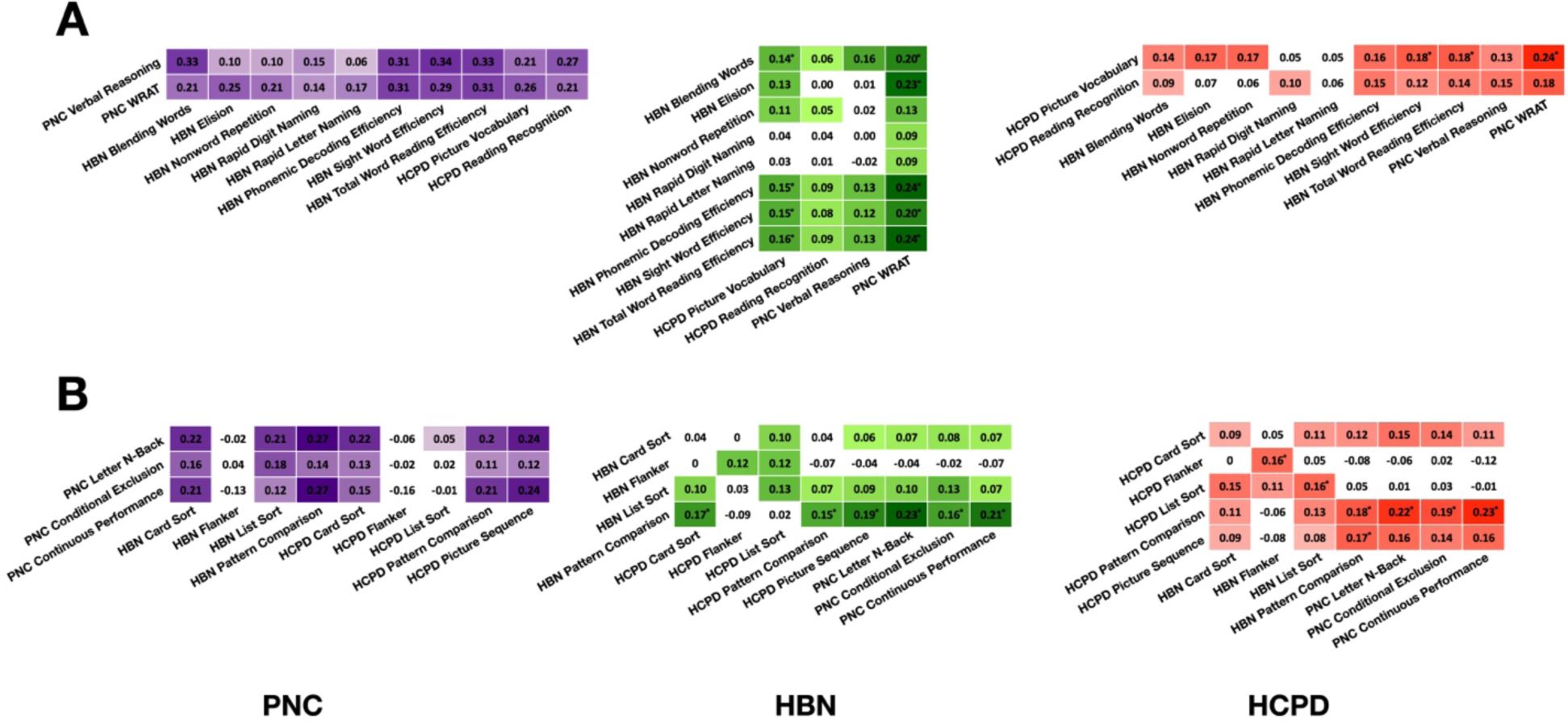
Cross-dataset predictions of individual measures. Models were trained on a single measure in one dataset (x-axis) and independently tested on each individual measure of the other dataset (y-axis) for language abilities (A) and executive function (B). Performance r values are shown for PNC (purple), HBN (green), and HCPD (red). Darker colors indicate higher prediction performances. White indicates non-significant performances. Asterisks indicate predictions greater than PCA-derived cross-dataset predictions.

Finally, we correlated within-dataset and cross-dataset performance. The ability of a measure to predict measures in another dataset did not correlate with its within-dataset performance (r=0.21, p=0.34). However, the ability of a measure to be predicted by measures in another dataset strongly correlated with within-dataset performance (r=0.72, p<0.001). These results indicate that a measure’s within-dataset performance estimates its predictability from other models, but not the predictive ability of its model on other measures.

## DISCUSSION

We used connectome-based predictive modeling to test the generalizability of neuroimaging predictive models across diverse dataset features. Predictions of language abilities and EF survived testing across three unharmonized, large-scale developmental samples. These results suggest reproducible associations that overcome individual dataset idiosyncrasies can be achieved with sample sizes (n=500-1000’s) below consortium-level magnitudes. Further, many models based on an individual measure of language or EF generalized to different language or EF measures. Interestingly, both PCA and individual measure results indicate that a model’s within-dataset performance estimates its predictability from other models but not the predictive ability of its model on other measures. Testing brain-behavior associations across diverse data remains necessary to strengthen the generalizability of findings beyond a particular dataset and assess applicability to real-world settings.

Our results highlight the potential of pooling neuroimaging data without harmonization. Notably, for the HCPD and HBN datasets, the best predictions were not from training and testing in the same dataset (i.e., cross-validation) but from external validation. This result suggests that training on diverse data may improve prediction in specific cases. Of course, strictly harmonized data collection efforts by consortiums remain essential (Casey et al., 2018; Sudlow et al., 2015). They maximize statistical power by minimizing unexplained variance (i.e., experimental noise). Nevertheless, harmonization is expensive and not always possible (Chow et al., 2023; Torres-Espín and Ferguson, 2022). It also prevents testing a model’s robustness to different experimental factors. Thus, testing on non-harmonized data is needed. While post-hoc harmonization (i.e., ComBat) is often applied in these studies, we avoided this step to test how brain-behavior associations can generalize without explicit harmonization (Chen et al., 2022; Yan et al., 2023). Using non-harmonized sources is a strength of neuroimaging predictive modeling. Recent work suggests that machine learning models predicting treatment outcomes from clinical data may fail when applied to unharmonized samples (Chekroud et al., 2024). Our results point to the potential value of incorporating neuroimaging data to improve generalization across unharmonized samples.

Though our models generalize well, lacking generalization is not inherently bad. A single model will not be appropriate in all cases. For example, models designed for adults likely should not work on infants and young children (Scheinost et al., 2023). Many brain-behavior associations may exhibit sex differences, where sex-specific models could be needed (Dhamala et al., 2023; Greene et al., 2018; Jiang et al., 2020; Yip et al., 2023). Further, evidence suggests that those who defy stereotypes (such as minoritized populations) could require different models (Greene et al., 2022). Rigorously testing a model on diverse data, regardless of whether it generalizes, produces valuable information. Null results motivate future studies to understand the lack of generalization and should be published (Munafò and Neill, 2016). As a field, we should encourage testing models on diverse data to understand the effects of dataset shift and if models generalize.

We employed state-of-the-field methodology to use as much data as possible. This approach includes using large sample sizes to create and externally validate models. In contrast to most studies using external validation, the sample sizes for external validation were of the same order as the training data (Rosenblatt et al., 2023; Yeung et al., 2022). In fact, given that two external datasets were used to validate each model, more data was used to test a model than train it. This approach ensured we had adequate power for external validation. In all cases, we had at least 80% power for effects as low as r=0.15. In addition to using large sample sizes, we also used several fMRI runs and multiple behavior measures for each individual. Combining fMRI and behavior data improves prediction likely by averaging out the idiosyncrasies of each data point and increasing reliability. These latent factors also allow diverse data types (i.e., different fMRI tasks and behavioral measures) to be used for prediction. Finally, we preserved participants without imaging data to derive principal components (e.g., using 6745 PNC and 1281 HBN participants) to increase the representation. These results follow the growing appreciation of large (i.e., many participants) and deep (i.e., many measures per participant) data (Gordon et al., 2017; Marek et al., 2022).

Statistical power remains a fundamental consideration in neuroimaging (Cremers et al., 2017). A rule of thumb is often desired (i.e., 1,000 participants are needed for an fMRI experiment). However, a simple answer is often insufficient given the complexities of relating neuroimaging data to behavior. There are too many modalities, behaviors, and analysis methods. Though, some generalities can be made. Our results demonstrate that predictive models can generalize across diverse, unharmonized data. These findings underscore the potential to employ neuroimaging models for predicting personalized outcomes and finding robust brain-behavior associations (Spisak et al., 2023). Of course, results will likely be case-specific. Language and EF exhibit large effect sizes for brain-behavior associations. Other behaviors and phenotypes, such as clinical symptoms, may need larger samples or improved methodology to create robust associations.

Executive function and language abilities are core cognitive processes that are critical for everyday functioning. Executive function supports manipulating information to plan, organize, and execute decisions towards goal-directed tasks (Cristofori et al., 2019; Diamond, 2013). Language abilities support the effective production and comprehension of communication toward meaningful interaction (Kidd et al., 2018). Cognitive deficits are associated with a range of psychiatric and developmental disorders (Millan et al., 2012; Zelazo, 2020). Achieving robust predictions of these constructs is meaningful for cognitive and clinical neuroscience (Barron et al., 2020; Boyle et al., 2023; Sui et al., 2020). However, the observed effect sizes are still smaller than necessary for real-world utility. Further, even if our models were actionable, ethical concerns related to their implementation in developmental populations exist (Scheinost et al., 2023). For example, false positives lead to unnecessary interventions, while false negatives divert resources from those who need them. Another consideration is model interpretability. Clinicians may be more hesitant to trust and integrate less interpretable models into their practice (Chekroud et al., 2021). The edges we observed contributing to language abilities and executive function predictions were distributed throughout the brain. It is difficult to pinpoint a single canonical network responsible for individual variation in performance (Kohoutová et al., 2020). However, these models align with recent literature that appreciates complex brain-wide networks rather than the simple networks often identified by traditional association studies (Dubois et al., 2018).

The strength of this study is the rigorous validation of the models. First, we used three large developmental datasets to maximize statistical power. Few large-scale neuroimaging studies incorporate any form of external validation (Rosenblatt et al., 2023; Yeung et al., 2022). In addition to internal cross-validation, each model was validated in two independent large-scale datasets. Future applications of brain-based predictive modeling methods must overcome demographics, imaging, and behavioral data differences. The three datasets exhibited substantial variability in participant demographics, geographic distribution, and clinical symptoms. Further, the notable lack of harmonization suggests that these models are not dependent upon specific study designs or measurement features. Thus, our results are highly generalizable and robust to dataset shift.

Several limitations exist. Using PCA on disparate behavioral measures may inadvertently remove some elements that make each measure unique. For example, unique components of EF include working memory, cognitive flexibility, and inhibitory control. Thus, latent measures from PCA might not represent these components but instead represent general cognition (Dyer and Kording, 2023). Similarly, we define language abilities broadly, including receptive language, expressive language, speech, and reading measures. These broad definitions may also explain the models’ lack of localization. More specific phenotypes will likely improve a model’s interpretability (Enkavi et al., 2019; Greene and Constable, 2023). We also see strong cross-dataset predictions for individual measures, so testing this hypothesis is plausible for future work. While our models generalized across various factors, all datasets were developmental samples from the United States. It is unclear if models would generalize to older individuals or those from non-western countries.

In conclusion, we show that brain-behavior associations generated from functional connectivity data can generalize over non-harmonized data. These results highlight that generalizable models can be achieved with datasets below consortium-level sample sizes and the potential of using non-harmonized data. Mimicking real-world dataset shifts in training and testing predictive models may accelerate their development into clinical tools and practice.

## METHODS

### Datasets

PNC participants were 1291 individuals ages 8-21 recruited from the greater Philadelphia, Pennsylvania area (Satterthwaite et al., 2016). Participants completed rest, emotion task, and n-back task fMRI runs (Satterthwaite et al., 2014). Measures of language abilities were the Penn Verbal Reasoning Task from the Penn Computerized Neurocognitive Battery (CNB) and the total standard score from the Wide Range Assessment Test (WRAT) Reading Subscale (Gur et al., 2010; Wilkinson and Robertson, 2006). Executive function measures were the Letter N-Back, Conditional Exclusion, and Continuous Performance tasks from the CNB.

HBN participants were 1110 individuals ages 6-17 recruited from the New York City, New York region (Alexander et al., 2017). Participants completed two rest fMRI runs as well as ‘Despicable Me’ and ‘The Present’ movie-watching scan sessions. Measures of language abilities were the Elision, Blending Words, Nonword Repetition, Rapid Digit Naming, and Rapid Letter Naming scaled scores from the Comprehensive Test of Phonological Processing (CTOPP-2) and the Phonemic Decoding Efficiency, Sight Word Efficiency, and Total Word Reading Efficiency scaled scores from the Test of Word Reading Efficiency (TOWRE-2) (Dickens et al., 2015; Tarar et al., 2015). Executive function measures were the Flanker Inhibitory Control and Attention, List Sorting Working Memory, Pattern Comparison Processing Speed, and Dimensional Change Card Sort age-corrected standard scores from the NIH Toolbox (Weintraub et al., 2013).

HCPD participants were 428 individuals ages 8-22 recruited from St. Louis, Missouri, Twin Cities, Minnesota, Boston, Massachusetts, and Los Angeles, California (Somerville et al., 2018). Participants completed rest fMRI runs (Harms et al., 2018). Measures of language abilities were the Picture Vocabulary and Oral Reading Recognition age-corrected standard scores from the NIH Toolbox. Executive function measures were the Flanker Inhibitory Control and Attention, List Sorting Working Memory, Pattern Comparison Processing Speed, Dimensional Change Card Sort, and Picture Sequence Memory age-corrected standard scores from the NIH Toolbox.

### Preprocessing

In all datasets, data were motion-corrected. Additional preprocessing steps were performed using BioImage Suite (Papademetris et al., 2006). This included regression of covariates of no interest from the functional data, including linear and quadratic drifts, mean cerebrospinal fluid signal, mean white matter signal, and mean global signal. Additional motion control was applied by regressing a 24-parameter motion model, which included six rigid body motion parameters, six temporal derivatives, and the square of these terms, from the data. Subsequently, we applied temporal smoothing with a Gaussian filter (approximate cutoff frequency=0.12 Hz) and gray matter masking, as defined in common space. Then, the Shen 268-node atlas was applied to parcellate the denoised data into 268 nodes (Shen et al., 2013). Finally, we generated functional connectivity matrices by correlating each node time series data pair and applying the Fisher transform. Data were excluded for poor data quality, missing nodes due to lack of full brain coverage, high motion (>0.2mm mean frame-wise motion), or missing behavioral/phenotypic data.

### Creating latent factors of language abilities and EF

A principal components analysis (PCA) combined language abilities and EF measures, respectively, for each dataset. Here, a single behavioral measurement represents a noisy approximation of the behavioral construct. Combining across multiple measures reduces this noise. To maintain separate train and test groups in PNC and HBN, each PCA was limited to participants who did not have usable neuroimaging data (n=6745 for PNC, n=1281 for HBN).

### Ridge regression Connectome-based Predictive Modeling

Based on ridge regression, we modify the original CPM framework to better suit the high-dimensional nature of connectivity data (Gao et al., 2019). Specifically, due to the positive semi-definite nature of a functional connectivity matrix, the edges are not independent. Ridge regression is more robust than OLS in this case. Instead of summing selected edges and fitting a one-dimensional OLS model, we directly fit a ridge regression model with training individuals using the selected edges from all the tasks and apply the model to testing individuals in the cross-validation framework. We trained a ridge regression model using 10-fold cross-validation for the within-dataset models. We used Pearson’s correlation and a feature selection threshold of p<0.05. When controlling for confounds, partial correlation was used for feature selection. The L2 regularization parameter λ parameter was chosen by an inner 10-fold cross-validation which uses only the training individuals. The largest λ value with a mean squared error (MSE) within one standard error of the minimum MSE is chosen. This cross-validation was repeated for 100 random divisions.

### Model performance

Within dataset prediction was evaluated with a cross-validated coefficient of determination (q^2^), and the median q^2^ for 100 random 10-fold divisions is reported, along with Pearson’s correlation (r) and mean square error (MSE) (Poldrack et al., 2020). To generate null distributions for significance testing, we randomly shuffled the correspondence between behavioral variables and connectivity matrices 1,000 times and re-ran the CPM analysis with the shuffled data. Based on these null distributions, the p-values for predictions were calculated as in prior work. Only a positive association between predicted and actual values indicates prediction above chance (with negative associations indicating a failure to predict), so one-tailed p-values are reported. Pearson’s correlation was tested between actual and predicted values to evaluate cross-dataset prediction.

### Model contribution

Predictive networks identified using CPM are complex and composed of multiple brain regions and networks. To quantify the contribution of each edge to a given predictive model, we calculated the *k*^*th*^ edge’s weight (labeled *W_k_*,) to the model as: *W_k_* = *abs*(*β^k^*)*std*(*E_k_*), where *std*(*E_k_*) represents the standard deviation of the *k^th^* edge, and *β^k^* represents the weight learned by CPM for the *k^th^* edge. To quantify the contribution of each node to a given predictive model, we calculated the *n^th^* node’s weight summed across all edges (labeled *W*_n_) to the model as: 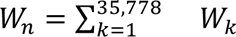, for all *k* edges connected to the *n^th^* node. Next, for the network level, *W_k_* was averaged over each edge within or between canonical functional networks.

### Virtual lesioning

CPM predictive networks are typically widespread and complex, so we conducted a virtual lesion analysis. For a CPM-based virtual lesion analysis, predictive networks can be set to zero to examine the degradation in predictive performance attributed to a virtual lesion of that network (Yip et al., 2020). We iteratively set each functional network to zero and examined how this impacted the model performance. We conducted this virtual lesion analysis for the canonical functional networks: medial frontal (MF), frontoparietal (FP), default mode (DMN), motor (MOT), visual I (VI), visual II (VII), visual association (VA), salience (SAL), subcortical (SC), and cerebellum (CBL).

### Data availability

Data are available through the Healthy Brain Network Dataset (https://data.healthybrainnetwork.org/main.php), the Human Connectome Project in Development Dataset (https://nda.nih.gov/), and the Philadelphia Neurodevelopmental Cohort Dataset (https://www.ncbi.nlm.nih.gov/projects/gap/cgi-bin/study.cgi?study_id=phs000607.v3.p2).

### Code availability

Preprocessing was carried out using Bioimage Suite, which is freely available: https://medicine.yale.edu/bioimaging/suite/. Code for the analyses is available at: https://github.com/brendan-adkinson/generalization/.

## REFERENCES

Adise, S., Ottino-Gonzalez, J., Goedde, L., Marshall, A.T., Kan, E., Rhee, K.E., Goran, M.I., Sowell, E.R., 2023. Variation in executive function relates to BMI increases in youth who were initially of a healthy weight in the ABCD Study. Obesity 31, 2809–2821. 10.1002/oby.23811

Alexander, L.M., Escalera, J., Ai, L., Andreotti, C., Febre, K., Mangone, A., Vega-Potler, N., Langer, N., Alexander, A., Kovacs, M., Litke, S., O’Hagan, B., Andersen, J., Bronstein, B., Bui, A., Bushey, M., Butler, H., Castagna, V., Camacho, N., Chan, E., Citera, D., Clucas, J., Cohen, S., Dufek, S., Eaves, M., Fradera, B., Gardner, J., Grant-Villegas, N., Green, G., Gregory, C., Hart, E., Harris, S., Horton, M., Kahn, D., Kabotyanski, K., Karmel, B., Kelly, S.P., Kleinman, K., Koo, B., Kramer, E., Lennon, E., Lord, C., Mantello, G., Margolis, A., Merikangas, K.R., Milham, J., Minniti, G., Neuhaus, R., Levine, A., Osman, Y., Parra, L.C., Pugh, K.R., Racanello, A., Restrepo, A., Saltzman, T., Septimus, B., Tobe, R., Waltz, R., Williams, A., Yeo, A., Castellanos, F.X., Klein, A., Paus, T., Leventhal, B.L., Craddock, R.C., Koplewicz, H.S., Milham, M.P., 2017. An open resource for transdiagnostic research in pediatric mental health and learning disorders. Sci. Data 4, 170181. 10.1038/sdata.2017.181

Barron, D.S., Gao, S., Dadashkarimi, J., Greene, A.S., Spann, M.N., Noble, S., Lake, E.M.R., Krystal, J.H., Constable, R.T., Scheinost, D., 2020. Transdiagnostic, Connectome-Based Prediction of Memory Constructs Across Psychiatric Disorders. Cereb. Cortex N. Y. NY 31, 2523–2533. 10.1093/cercor/bhaa371

Boyle, R., Connaughton, M., McGlinchey, E., Knight, S.P., De Looze, C., Carey, D., Stern, Y., Robertson, I.H., Kenny, R.A., Whelan, R., 2023. Connectome-based predictive modelling of cognitive reserve using task-based functional connectivity. Eur. J. Neurosci. 57, 490–510. 10.1111/ejn.15896

Casey, B.J., 2023. Executive functions in the brain, development and social context: Early contributions by neuroscientist, Adele Diamond. Dev. Cogn. Neurosci. 62, 101272. 10.1016/j.dcn.2023.101272

Casey, B.J., Cannonier, T., Conley, M.I., Cohen, A.O., Barch, D.M., Heitzeg, M.M., Soules, M.E., Teslovich, T., Dellarco, D.V., Garavan, H., Orr, C.A., Wager, T.D., Banich, M.T., Speer, N.K., Sutherland, M.T., Riedel, M.C., Dick, A.S., Bjork, J.M., Thomas, K.M., Chaarani, B., Mejia, M.H., Hagler, D.J., Daniela Cornejo, M., Sicat, C.S., Harms, M.P., Dosenbach, N.U.F., Rosenberg, M., Earl, E., Bartsch, H., Watts, R., Polimeni, J.R., Kuperman, J.M., Fair, D.A., Dale, A.M., ABCD Imaging Acquisition Workgroup, 2018. The Adolescent Brain Cognitive Development (ABCD) study: Imaging acquisition across 21 sites. Dev. Cogn. Neurosci. 32, 43–54. 10.1016/j.dcn.2018.03.001

Chekroud, A.M., Bondar, J., Delgadillo, J., Doherty, G., Wasil, A., Fokkema, M., Cohen, Z., Belgrave, D., DeRubeis, R., Iniesta, R., Dwyer, D., Choi, K., 2021. The promise of machine learning in predicting treatment outcomes in psychiatry. World Psychiatry 20, 154–170. 10.1002/wps.20882

Chekroud, A.M., Hawrilenko, M., Loho, H., Bondar, J., Gueorguieva, R., Hasan, A., Kambeitz, J., Corlett, P.R., Koutsouleris, N., Krumholz, H.M., Krystal, J.H., Paulus, M., 2024. Illusory generalizability of clinical prediction models. Science 383, 164–167. 10.1126/science.adg8538

Chen, A.A., Luo, C., Chen, Y., Shinohara, R.T., Shou, H., Alzheimer’s Disease Neuroimaging Initiative, 2022. Privacy-preserving harmonization via distributed ComBat. NeuroImage 248, 118822. 10.1016/j.neuroimage.2021.118822

Chow, S.-M., Nahum-Shani, I., Baker, J.T., Spruijt-Metz, D., Allen, N.B., Auerbach, R.P., Dunton, G.F., Friedman, N.P., Intille, S.S., Klasnja, P., Marlin, B., Nock, M.K., Rauch, S.L., Pavel, M., Vrieze, S., Wetter, D.W., Kleiman, E.M., Brick, T.R., Perry, H., Wolff-Hughes, D.L., Intensive Longitudinal Health Behavior Network (ILHBN), 2023. The ILHBN: challenges, opportunities, and solutions from harmonizing data under heterogeneous study designs, target populations, and measurement protocols. Transl. Behav. Med. 13, 7–16. 10.1093/tbm/ibac069

Cremers, H.R., Wager, T.D., Yarkoni, T., 2017. The relation between statistical power and inference in fMRI. PloS One 12, e0184923. 10.1371/journal.pone.0184923

Cristofori, I., Cohen-Zimerman, S., Grafman, J., 2019. Chapter 11 - Executive functions, in: D’Esposito, M., Grafman, J.H. (Eds.), Handbook of Clinical Neurology, The Frontal Lobes. Elsevier, pp. 197–219. 10.1016/B978-0-12-804281-6.00011-2

Dhamala, E., Rong Ooi, L.Q., Chen, J., Ricard, J.A., Berkeley, E., Chopra, S., Qu, Y., Zhang, X.-H., Lawhead, C., Yeo, B.T.T., Holmes, A.J., 2023. Brain-Based Predictions of Psychiatric Illness-Linked Behaviors Across the Sexes. Biol. Psychiatry 94, 479–491. 10.1016/j.biopsych.2023.03.025

Diamond, A., 2013. Executive Functions. Annu. Rev. Psychol. 64, 135. 10.1146/annurev-psych-113011-143750

Dickens, R.H., Meisinger, E.B., Tarar, J.M., 2015. Test Review: Comprehensive Test of Phonological Processing–2nd ed. (CTOPP-2) by Wagner, R. K., Torgesen, J. K., Rashotte, C. A., & Pearson, N. A. Can. J. Sch. Psychol. 30, 155–162. 10.1177/0829573514563280

Dockès, J., Varoquaux, G., Poline, J.-B., 2021. Preventing dataset shift from breaking machine-learning biomarkers. GigaScience 10, giab055. 10.1093/gigascience/giab055

Dubois, J., Adolphs, R., 2016. Building a Science of Individual Differences from fMRI. Trends Cogn. Sci. 20, 425–443. 10.1016/j.tics.2016.03.014

Dubois, J., Galdi, P., Paul, L.K., Adolphs, R., 2018. A distributed brain network predicts general intelligence from resting-state human neuroimaging data. Philos. Trans. R. Soc. B Biol. Sci. 373, 20170284. 10.1098/rstb.2017.0284

Dyer, E.L., Kording, K., 2023. Why the simplest explanation isn’t always the best. Proc. Natl. Acad. Sci. 120, e2319169120. 10.1073/pnas.2319169120

Elliott, M.L., Knodt, A.R., Cooke, M., Kim, M.J., Melzer, T.R., Keenan, R., Ireland, D., Ramrakha, S., Poulton, R., Caspi, A., Moffitt, T.E., Hariri, A.R., 2019. General functional connectivity: Shared features of resting-state and task fMRI drive reliable and heritable individual differences in functional brain networks. Neuroimage 189, 516–532. 10.1016/j.neuroimage.2019.01.068

Enkavi, A.Z., Eisenberg, I.W., Bissett, P.G., Mazza, G.L., MacKinnon, D.P., Marsch, L.A., Poldrack, R.A., 2019. Large-scale analysis of test–retest reliabilities of self-regulation measures. Proc. Natl. Acad. Sci. 116, 5472–5477. 10.1073/pnas.1818430116

Gao, S., Greene, A.S., Constable, R.T., Scheinost, D., 2019. Combining multiple connectomes improves predictive modeling of phenotypic measures. NeuroImage 201, 116038. 10.1016/j.neuroimage.2019.116038

Genon, S., Eickhoff, S.B., Kharabian, S., 2022. Linking interindividual variability in brain structure to behaviour. Nat. Rev. Neurosci. 23, 307–318. 10.1038/s41583-022-00584-7

Godfrey, K.J., Espenhahn, S., Stokoe, M., McMorris, C., Murias, K., McCrimmon, A., Harris, A.D., Bray, S., 2022. Autism interest intensity in early childhood associates with executive functioning but not reward sensitivity or anxiety symptoms. Autism 26, 1723–1736. 10.1177/13623613211064372

Gordon, E.M., Laumann, T.O., Gilmore, A.W., Newbold, D.J., Greene, D.J., Berg, J.J., Ortega, M., Hoyt-Drazen, C., Gratton, C., Sun, H., Hampton, J.M., Coalson, R.S., Nguyen, A.L., McDermott, K.B., Shimony, J.S., Snyder, A.Z., Schlaggar, B.L., Petersen, S.E., Nelson, S.M., Dosenbach, N.U.F., 2017. Precision Functional Mapping of Individual Human Brains. Neuron 95, 791–807.e7. 10.1016/j.neuron.2017.07.011

Greene, A.S., Constable, R.T., 2023. Clinical Promise of Brain-Phenotype Modeling: A Review. JAMA Psychiatry 80, 848. 10.1001/jamapsychiatry.2023.1419

Greene, A.S., Gao, S., Scheinost, D., Constable, R.T., 2018. Task-induced brain state manipulation improves prediction of individual traits. Nat. Commun. 9, 2807. 10.1038/s41467-018-04920-3

Greene, A.S., Shen, X., Noble, S., Horien, C., Hahn, C.A., Arora, J., Tokoglu, F., Spann, M.N., Carrión, C.I., Barron, D.S., Sanacora, G., Srihari, V.H., Woods, S.W., Scheinost, D., Constable, R.T., 2022. Brain-phenotype models fail for individuals who defy sample stereotypes. Nature 609, 109–118. 10.1038/s41586-022-05118-w

Gur, R.C., Richard, J., Hughett, P., Calkins, M.E., Macy, L., Bilker, W.B., Brensinger, C., Gur, R.E., 2010. A cognitive neuroscience based computerized battery for efficient measurement of individual differences: Standardization and initial construct validation. J. Neurosci. Methods 187, 254–262. 10.1016/j.jneumeth.2009.11.017

Harms, M.P., Somerville, L.H., Ances, B.M., Andersson, J., Barch, D.M., Bastiani, M., Bookheimer, S.Y., Brown, T.B., Buckner, R.L., Burgess, G.C., Coalson, T.S., Chappell, M.A., Dapretto, M., Douaud, G., Fischl, B., Glasser, M.F., Greve, D.N., Hodge, C., Jamison, K.W., Jbabdi, S., Kandala, S., Li, X., Mair, R.W., Mangia, S., Marcus, D., Mascali, D., Moeller, S., Nichols, T.E., Robinson, E.C., Salat, D.H., Smith, S.M., Sotiropoulos, S.N., Terpstra, M., Thomas, K.M., Tisdall, M.D., Ugurbil, K., van der Kouwe, A., Woods, R.P., Zöllei, L., Van Essen, D.C., Yacoub, E., 2018. Extending the Human Connectome Project across ages: Imaging protocols for the Lifespan Development and Aging projects. NeuroImage 183, 972–984. 10.1016/j.neuroimage.2018.09.060

Jiang, R., Calhoun, V.D., Fan, L., Zuo, N., Jung, R., Qi, S., Lin, D., Li, J., Zhuo, C., Song, M., Fu, Z., Jiang, T., Sui, J., 2020. Gender Differences in Connectome-based Predictions of Individualized Intelligence Quotient and Sub-domain Scores. Cereb. Cortex N. Y. N 1991 30, 888–900. 10.1093/cercor/bhz134

Jiang, R., Woo, C.-W., Qi, S., Wu, J., Sui, J., 2022. Interpreting Brain Biomarkers: Challenges and solutions in interpreting machine learning-based predictive neuroimaging. IEEE Signal Process. Mag. 39, 107–118. 10.1109/MSP.2022.3155951

Johnson, K.B., Wei, W., Weeraratne, D., Frisse, M.E., Misulis, K., Rhee, K., Zhao, J., Snowdon, J.L., 2021. Precision Medicine, AI, and the Future of Personalized Health Care. Clin. Transl. Sci. 14, 86–93. 10.1111/cts.12884

Kidd, E., Donnelly, S., Christiansen, M.H., 2018. Individual Differences in Language Acquisition and Processing. Trends Cogn. Sci. 22, 154–169. 10.1016/j.tics.2017.11.006

Klapwijk, E.T., van den Bos, W., Tamnes, C.K., Raschle, N.M., Mills, K.L., 2020.Opportunities for increased reproducibility and replicability of developmental neuroimaging. Dev. Cogn. Neurosci. 47, 100902. 10.1016/j.dcn.2020.100902

Kohoutová, L., Heo, J., Cha, S., Lee, S., Moon, T., Wager, T.D., Woo, C.-W., 2020. Toward a unified framework for interpreting machine-learning models in neuroimaging. Nat. Protoc. 15, 1399–1435. 10.1038/s41596-019-0289-5

Liu, S., Abdellaoui, A., Verweij, K.J.H., van Wingen, G.A., 2023. Replicable brain– phenotype associations require large-scale neuroimaging data. Nat. Hum. Behav. 7, 1344–1356. 10.1038/s41562-023-01642-5

Marek, S., Tervo-Clemmens, B., Calabro, F.J., Montez, D.F., Kay, B.P., Hatoum, A.S., Donohue, M.R., Foran, W., Miller, R.L., Hendrickson, T.J., Malone, S.M., Kandala, S., Feczko, E., Miranda-Dominguez, O., Graham, A.M., Earl, E.A., Perrone, A.J., Cordova, M., Doyle, O., Moore, L.A., Conan, G.M., Uriarte, J., Snider, K., Lynch, B.J., Wilgenbusch, J.C., Pengo, T., Tam, A., Chen, J., Newbold, D.J., Zheng, A., Seider, N.A., Van, A.N., Metoki, A., Chauvin, R.J., Laumann, T.O., Greene, D.J., Petersen, S.E., Garavan, H., Thompson, W.K., Nichols, T.E., Yeo, B.T.T., Barch, D.M., Luna, B., Fair, D.A., Dosenbach, N.U.F., 2022. Reproducible brain-wide association studies require thousands of individuals. Nature 603, 654–660. 10.1038/s41586-022-04492-9

Millan, M.J., Agid, Y., Brüne, M., Bullmore, E.T., Carter, C.S., Clayton, N.S., Connor, R., Davis, S., Deakin, B., DeRubeis, R.J., Dubois, B., Geyer, M.A., Goodwin, G.M., Gorwood, P., Jay, T.M., Joëls, M., Mansuy, I.M., Meyer-Lindenberg, A., Murphy, D., Rolls, E., Saletu, B., Spedding, M., Sweeney, J., Whittington, M., Young, L.J., 2012. Cognitive dysfunction in psychiatric disorders: characteristics, causes and the quest for improved therapy. Nat. Rev. Drug Discov. 11, 141–168. 10.1038/nrd3628

Munafò, M., Neill, J., 2016. Null is beautiful: On the importance of publishing null results. J. Psychopharmacol. (Oxf.) 30, 585–585. 10.1177/0269881116638813

Noble, S., Scheinost, D., Constable, R.T., 2020. Cluster failure or power failure? Evaluating sensitivity in cluster-level inference. NeuroImage 209, 116468. 10.1016/j.neuroimage.2019.116468

Papademetris, X., Jackowski, M.P., Rajeevan, N., DiStasio, M., Okuda, H., Constable, R.T., Staib, L.H., 2006. BioImage Suite: An integrated medical image analysis suite: An update. Insight J. 2006, 209.

Poldrack, R.A., Huckins, G., Varoquaux, G., 2020. Establishment of Best Practices for Evidence for Prediction: A Review. JAMA Psychiatry 77, 534–540. 10.1001/jamapsychiatry.2019.3671

Qi, T., Schaadt, G., Friederici, A.D., 2021. Associated functional network development and language abilities in children. NeuroImage 242, 118452. 10.1016/j.neuroimage.2021.118452

Rosenberg, M.D., Finn, E.S., 2022. How to establish robust brain–behavior relationships without thousands of individuals. Nat. Neurosci. 25, 835–837. 10.1038/s41593-022-01110-9

Rosenblatt, M., Tejavibulya, L., Camp, C.C., Jiang, R., Westwater, M.L., Noble, S., Scheinost, D., 2023. Power and reproducibility in the external validation of brain-phenotype predictions. BioRxiv Prepr. Serv. Biol. 2023.10.25.563971. 10.1101/2023.10.25.563971

Satterthwaite, T.D., Connolly, J.J., Ruparel, K., Calkins, M.E., Jackson, C., Elliott, M.A., Roalf, D.R., Hopson, R., Prabhakaran, K., Behr, M., Qiu, H., Mentch, F.D., Chiavacci, R., Sleiman, P.M.A., Gur, R.C., Hakonarson, H., Gur, R.E., 2016. The Philadelphia Neurodevelopmental Cohort: A publicly available resource for the study of normal and abnormal brain development in youth. NeuroImage 124, 1115–1119. 10.1016/j.neuroimage.2015.03.056

Satterthwaite, T.D., Elliott, M.A., Ruparel, K., Loughead, J., Prabhakaran, K., Calkins, M.E., Hopson, R., Jackson, C., Keefe, J., Riley, M., Mentch, F.D., Sleiman, P., Verma, R., Davatzikos, C., Hakonarson, H., Gur, R.C., Gur, R.E., 2014. Neuroimaging of the Philadelphia neurodevelopmental cohort. NeuroImage 86, 544–553. 10.1016/j.neuroimage.2013.07.064

Scheinost, D., Pollatou, A., Dufford, A.J., Jiang, R., Farruggia, M.C., Rosenblatt, M., Peterson, H., Rodriguez, R.X., Dadashkarimi, J., Liang, Q., Dai, W., Foster, M.L., Camp, C.C., Tejavibulya, L., Adkinson, B.D., Sun, H., Ye, J., Cheng, Q., Spann, M.N., Rolison, M., Noble, S., Westwater, M.L., 2023. Machine Learning and Prediction in Fetal, Infant, and Toddler Neuroimaging: A Review and Primer. Biol. Psychiatry 93, 893–904. 10.1016/j.biopsych.2022.10.014

Shen, X., Finn, E.S., Scheinost, D., Rosenberg, M.D., Chun, M.M., Papademetris, X., Constable, R.T., 2017. Using connectome-based predictive modeling to predict individual behavior from brain connectivity. Nat. Protoc. 12, 506–518. 10.1038/nprot.2016.178

Shen, X., Tokoglu, F., Papademetris, X., Constable, R.T., 2013. Groupwise whole-brain parcellation from resting-state fMRI data for network node identification. NeuroImage 82, 403–415. 10.1016/j.neuroimage.2013.05.081

Somerville, L.H., Bookheimer, S.Y., Buckner, R.L., Burgess, G.C., Curtiss, S.W., Dapretto, M., Stine Elam, J., Gaffrey, M.S., Harms, M.P., Hodge, C., Kandala, S., Kastman, E.K., Nichols, T.E., Schlaggar, B.L., Smith, S.M., Thomas, K.M., Yacoub, E., Van Essen, D.C., Barch, D.M., 2018. The Lifespan Human Connectome Project in Development: A large-scale study of brain connectivity development in 5–21 year olds. NeuroImage 183, 456–468. 10.1016/j.neuroimage.2018.08.050

Spisak, T., Bingel, U., Wager, T.D., 2023. Multivariate BWAS can be replicable with moderate sample sizes. Nature 615, E4–E7. 10.1038/s41586-023-05745-x

Sudlow, C., Gallacher, J., Allen, N., Beral, V., Burton, P., Danesh, J., Downey, P., Elliott, P., Green, J., Landray, M., Liu, B., Matthews, P., Ong, G., Pell, J., Silman, A., Young, A., Sprosen, T., Peakman, T., Collins, R., 2015. UK biobank: an open access resource for identifying the causes of a wide range of complex diseases of middle and old age. PLoS Med. 12, e1001779. 10.1371/journal.pmed.1001779

Sui, J., Jiang, R., Bustillo, J., Calhoun, V., 2020. Neuroimaging-based Individualized Prediction of Cognition and Behavior for Mental Disorders and Health: Methods and Promises. Biol. Psychiatry 88, 818–828. 10.1016/j.biopsych.2020.02.016

Tarar, J.M., Meisinger, E.B., Dickens, R.H., 2015. Test Review: Test of Word Reading Efficiency–Second Edition (TOWRE-2) by Torgesen, J. K., Wagner, R. K., & Rashotte, C. A. Can. J. Sch. Psychol. 30, 320–326. 10.1177/0829573515594334

Torres-Espín, A., Ferguson, A., 2022. Harmonization-Information Trade-Offs for Sharing Individual Participant Data in Biomedicine. Harv. Data Sci. Rev. 4. 10.1162/99608f92.a9717b34

Weintraub, S., Dikmen, S.S., Heaton, R.K., Tulsky, D.S., Zelazo, P.D., Bauer, P.J., Carlozzi, N.E., Slotkin, J., Blitz, D., Wallner-Allen, K., Fox, N.A., Beaumont, J.L., Mungas, D., Nowinski, C.J., Richler, J., Deocampo, J.A., Anderson, J.E., Manly, J.J., Borosh, B., Havlik, R., Conway, K., Edwards, E., Freund, L., King, J.W., Moy, C., Witt, E., Gershon, R.C., 2013. Cognition assessment using the NIH Toolbox. Neurology 80, S54–S64. 10.1212/WNL.0b013e3182872ded

Wilkinson, G.S., Robertson, G.J., 2006. Wide Range Achievement Test 4. 10.1037/t27160-000

Woo, C.-W., Chang, L.J., Lindquist, M.A., Wager, T.D., 2017. Building better biomarkers: brain models in translational neuroimaging. Nat. Neurosci. 20, 365–377. 10.1038/nn.4478

Yan, W., Fu, Z., Jiang, R., Sui, J., Calhoun, V.D., 2023. Maximum Classifier Discrepancy Generative Adversarial Network for Jointly Harmonizing Scanner Effects and Improving Reproducibility of Downstream Tasks. IEEE Trans. Biomed. Eng. 1–9. 10.1109/TBME.2023.3330087

Yarkoni, T., 2009. Big Correlations in Little Studies: Inflated fMRI Correlations Reflect Low Statistical Power—Commentary on Vul et al. (2009). Perspect. Psychol. Sci. 4, 294–298. 10.1111/j.1745-6924.2009.01127.x

Yarkoni, T., Westfall, J., 2017. Choosing prediction over explanation in psychology: Lessons from machine learning. Perspect. Psychol. Sci. J. Assoc. Psychol. Sci. 12, 1100–1122. 10.1177/1745691617693393

Yeung, A.W.K., More, S., Wu, J., Eickhoff, S.B., 2022. Reporting details of neuroimaging studies on individual traits prediction: A literature survey. NeuroImage 256, 119275. 10.1016/j.neuroimage.2022.119275

Yip, S.W., Kiluk, B., Scheinost, D., 2020. Toward Addiction Prediction: An Overview of Cross-Validated Predictive Modeling Findings and Considerations for Future Neuroimaging Research. Biol. Psychiatry Cogn. Neurosci. Neuroimaging, Understanding the Nature and Treatment of Psychopathology: Letting the Data Guide the Way 5, 748–758. 10.1016/j.bpsc.2019.11.001

Yip, S.W., Lichenstein, S.D., Liang, Q., Chaarani, B., Dager, A., Pearlson, G., Banaschewski, T., Bokde, A.L.W., Desrivières, S., Flor, H., Grigis, A., Gowland, P., Heinz, A., Brühl, R., Martinot, J.-L., Martinot, M.-L.P., Artiges, E., Nees, F., Orfanos, D.P., Paus, T., Poustka, L., Hohmann, S., Millenet, S., Fröhner, J.H., Smolka, M.N., Vaidya, N., Walter, H., Whelan, R., Schumann, G., Garavan, H., 2023. Brain Networks and Adolescent Alcohol Use. JAMA Psychiatry 80, 1131–1141. 10.1001/jamapsychiatry.2023.2949

Zelazo, P.D., 2020. Executive Function and Psychopathology: A Neurodevelopmental Perspective. Annu. Rev. Clin. Psychol. 16, 431–454. 10.1146/annurev-clinpsy-072319-024242

