## Supplement for "Brain-phenotype predictions can survive across diverse real-world data"

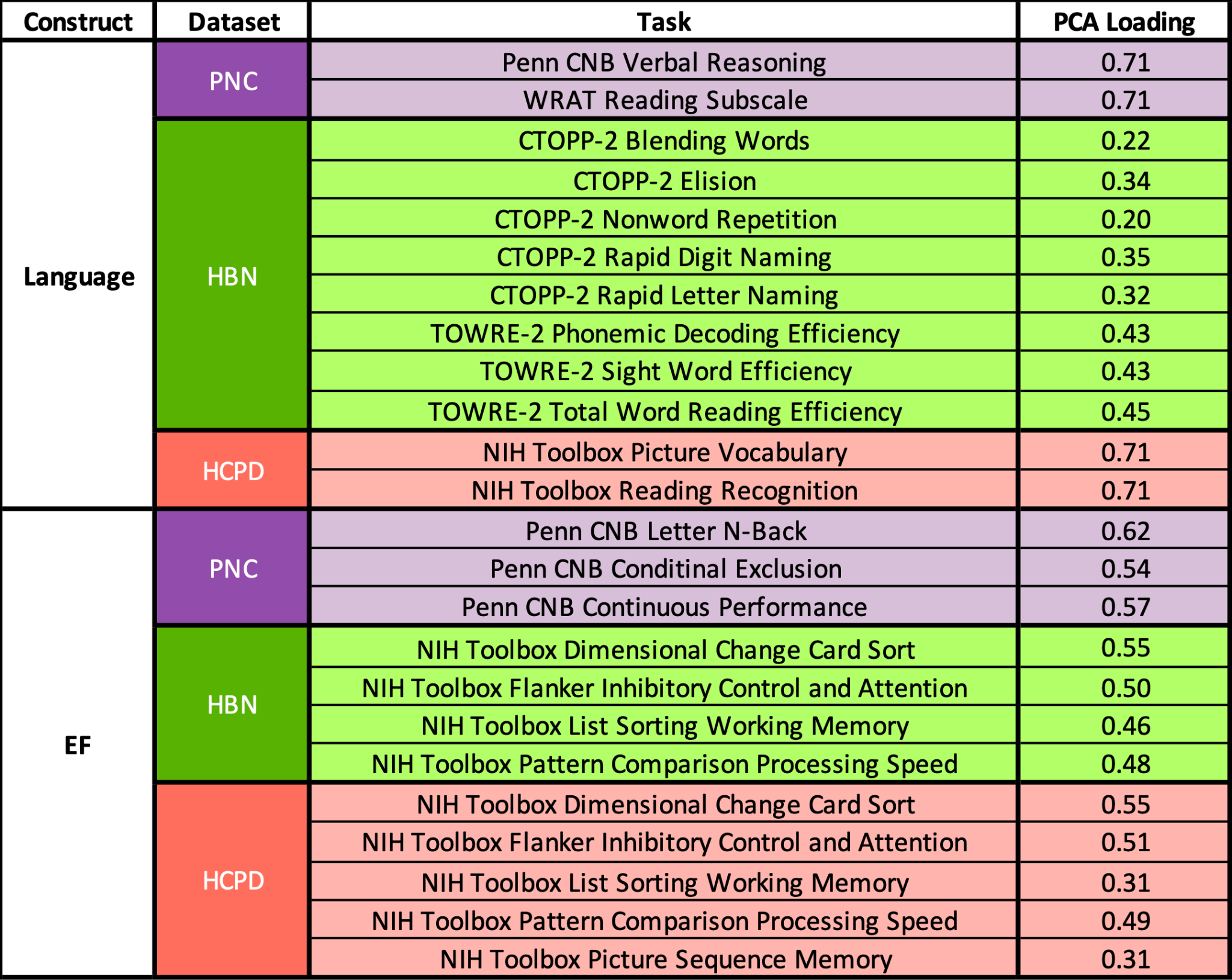

**Table S1. Individual Behavioral Tasks and Latent Factor Loadings.** The PNC (purple), HBN (green), and HCPD (red) datasets each used different behavioral tasks to assess language abilities (top) and executive function (bottom). The loadings of each measure to the PCA-derived “latent” factor for that dataset are presented in the final column.

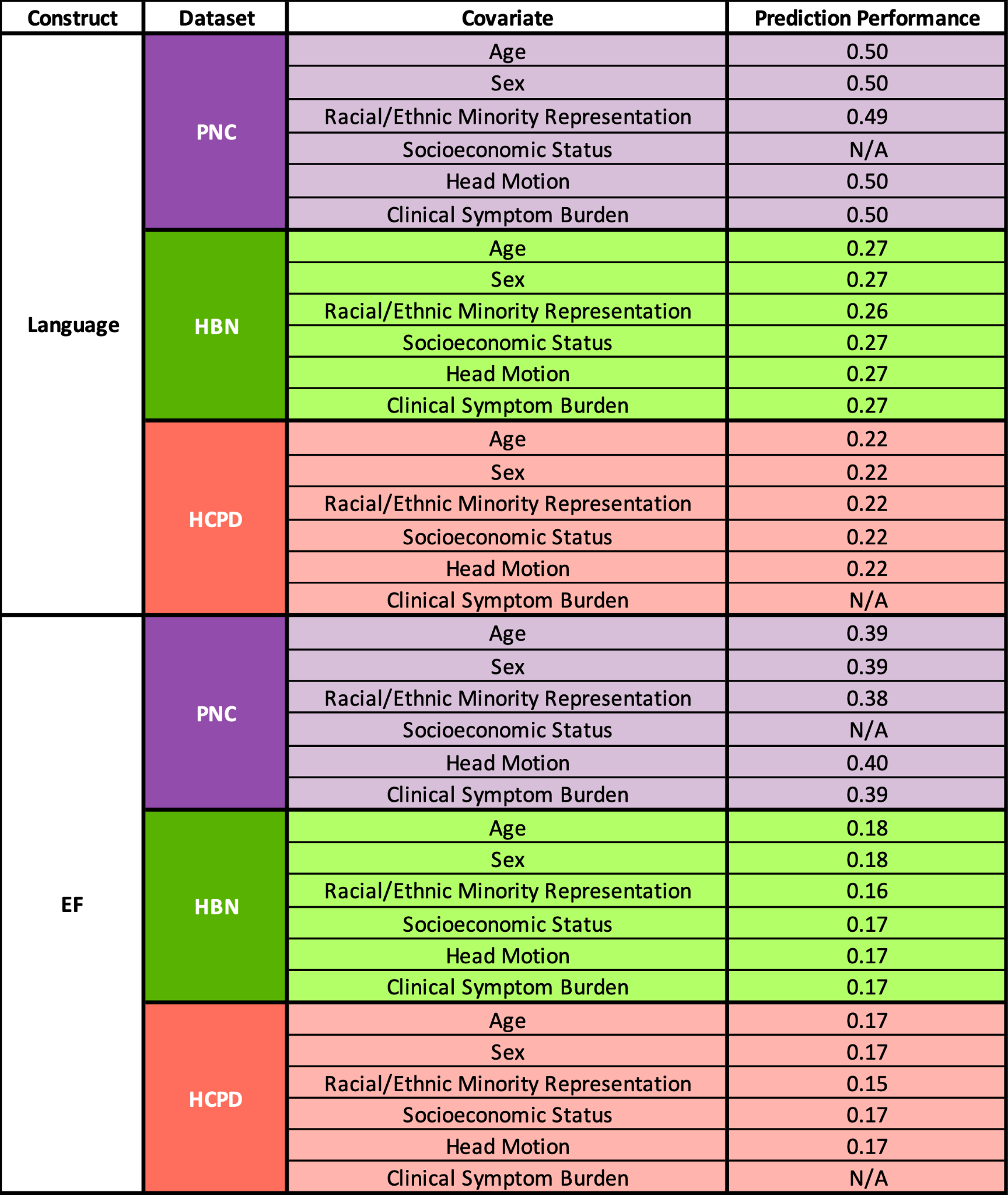

**Table S2. Model performances with addition of covariates.** Predictive models exhibited similar performances with the addition of covariates during feature selection for both language abilities (top) and executive function (bottom) across PNC (purple), HBN (green), and HCPD (red). Variables of interest were age, sex, racial/ethnic minority representation, socioeconomic status, head motion, and clinical symptom burdens.

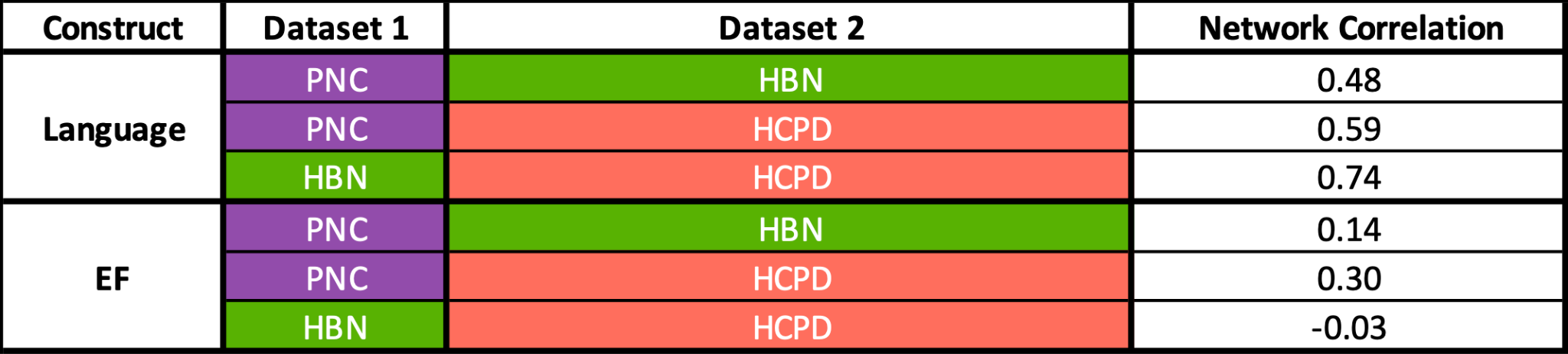

**Table S3. Comparison of Network-Level Brain Features Underlying Predictions.** For both language abilities (top) and executive function (bottom), we assessed the similarity of the networks underlying predictions between datasets (PNC: purple; HBN: green; HCPD: red). All edgewise regression coefficients were normalized by the standard deviation of edges and summed for each canonical brain network.

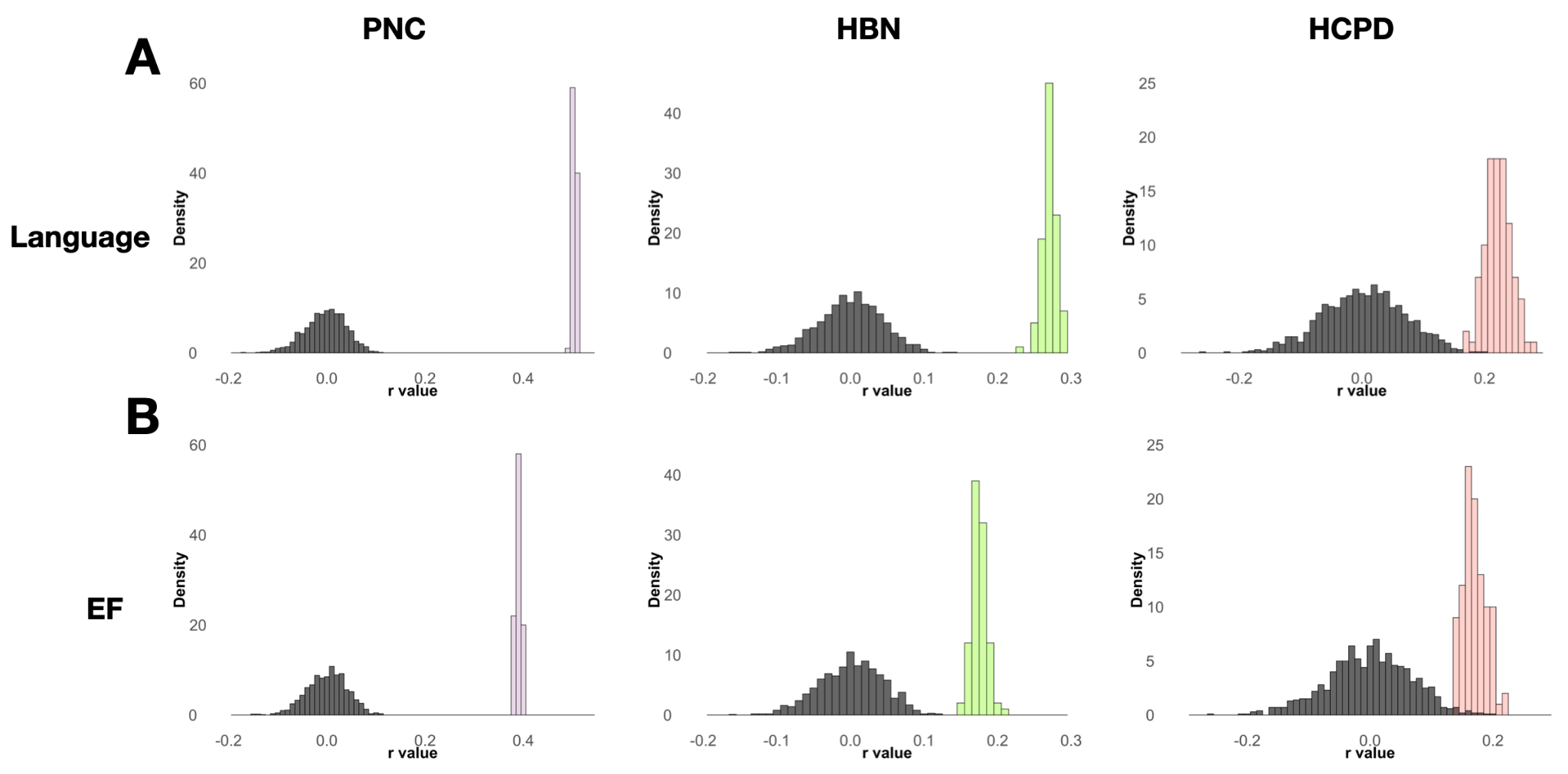

**Figure S1. Permutation Testing for Within-Dataset Predictions.** For within-dataset language abilities (A) and executive function (B) predictions, 100 iterations of 10-fold cross-validation were performed (PNC: purple; HBN: green; HCPD: red). Significance was assessed using permutation testing with 1000 iterations of randomly shuffled behavioral data labels (gray).

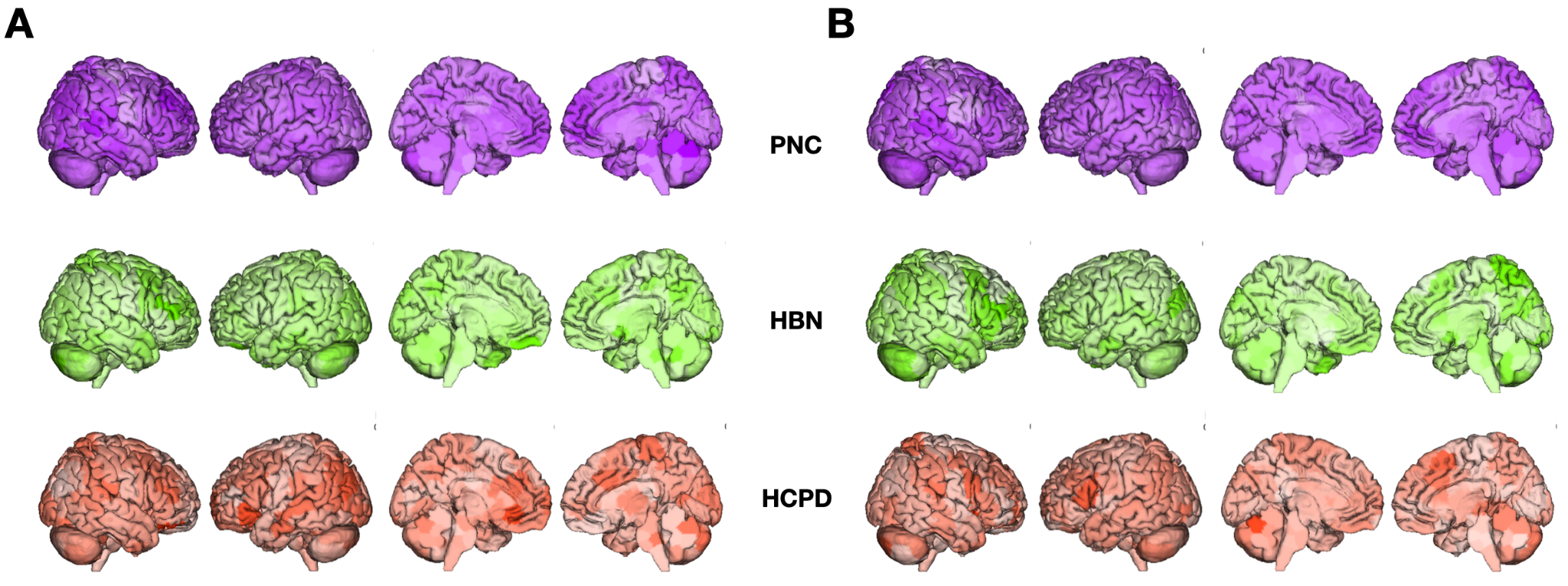

**Figure S2. Node-level contributions to language abilities and executive function predictions.** Node contributions are the sum of all edgewise ridge regression coefficients normalized by the standard deviation of edges across participants for a given node to predictions of language abilities (A) and executive function (B). PNC (top, purple), HBN (middle, green), and HCPD (bottom, red).

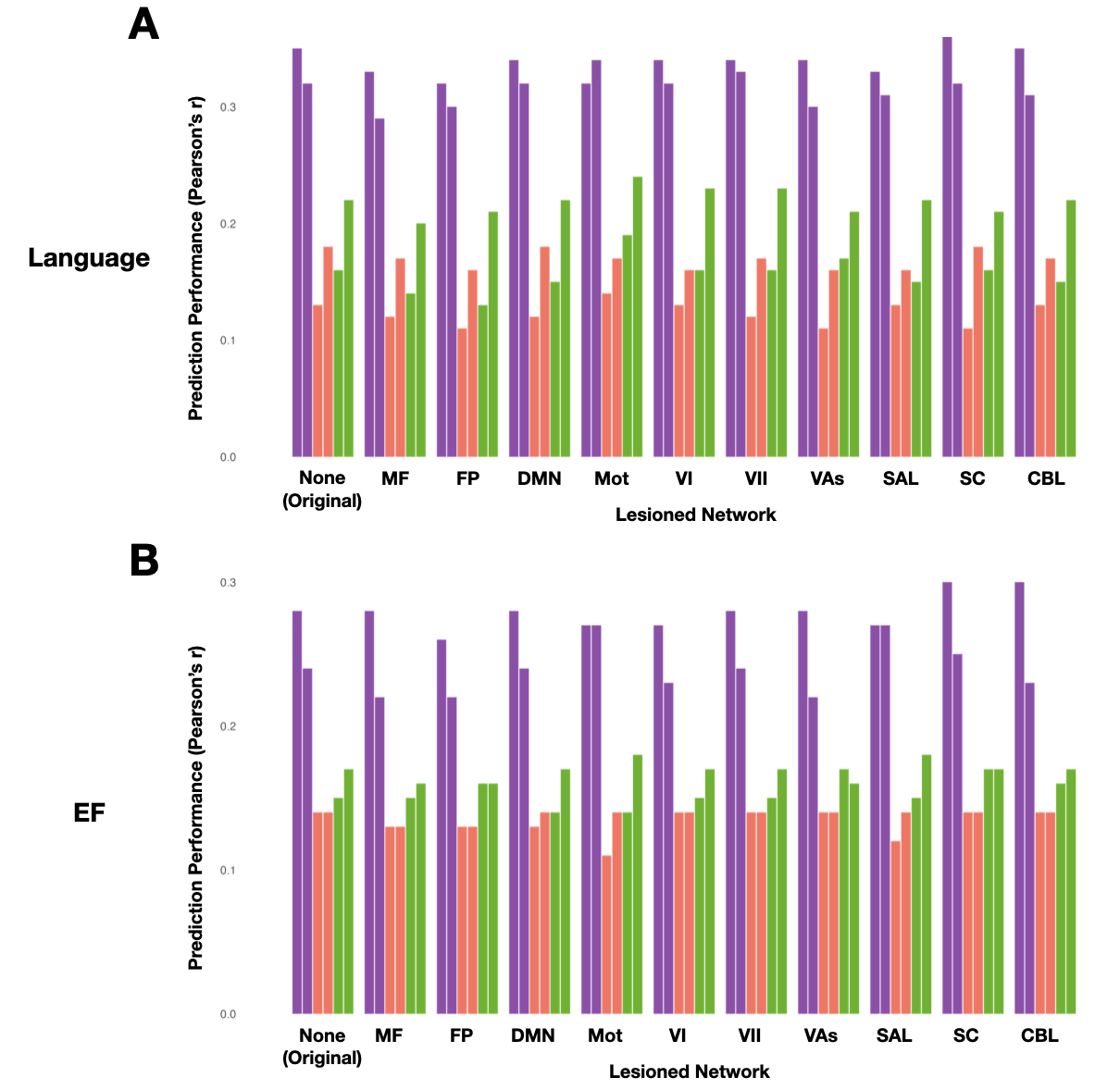

**Figure S3. Cross-Dataset Prediction Performances After Virtual Network Lesioning.** The impact of eliminating contributions of edges associated with given canonical networks on cross-dataset language abilities (A) and executive function (B) prediction performances. Colors indicate the dataset in which models were tested (PNC: purple; HBN: green; HCPD: red). Network Labels: MF, medial frontal; FP, frontoparietal; DMN, default mode; Mot, motor cortex; VI, visual A; VII, visual B; VAs, visual association; SAL, salience; SC, subcortical; CBL, cerebellum.
